# Database size positively correlates with the loss of species-level taxonomic resolution for the 16S rRNA and other prokaryotic marker genes

**DOI:** 10.1101/2023.12.13.571439

**Authors:** Seth Commichaux, Tu Luan, Harihara Subrahmaniam Muralidharan, Mihai Pop

**Affiliations:** Center for Food Safety and Nutrition, Food and Drug Administration, Laurel, MD 20708; Department of Computer Science, University of Maryland, College Park, MD 20742; Center for Bioinformatics and Computational Biology, University of Maryland, College Park, MD 20742

## Abstract

For decades, the 16S rRNA gene has been used to taxonomically classify prokaryotic species and to taxonomically profile microbial communities. The 16S rRNA gene has been criticized for being too conserved to differentiate between distinct species. We argue that the inability to differentiate between species is not a unique feature of the 16S rRNA gene. Rather, we observe the gradual loss of species-level resolution for other marker genes as the number of gene sequences increases in reference databases. We demonstrate this effect through the analysis of three commonly used databases of nearly-universal prokaryotic marker genes: the SILVA 16S rRNA gene database, the Genome Taxonomy Database (GTDB), and a set of 40 taxonomically-informative single-copy genes. Our results reflect a more fundamental property of the taxonomies themselves and have broad implications for bioinformatic analyses beyond taxonomic classification. Effective solutions for fine-level taxonomic classification require a more precise, and operationally-relevant, definition of the taxonomic labels being sought, and the use of combinations of genomic markers in the classification process.

**Importance:** The use of reference databases for assigning taxonomic labels to genomic and metagenomic sequences is a fundamental bioinformatic task in the characterization of microbial communities. The increasing accessibility of high throughput sequencing has led to a rapid increase in the size and number of sequences in databases. This has been beneficial for improving our understanding of the global microbial genetic diversity. However, there is evidence that as the microbial diversity is more densely sampled, increasingly longer genomic segments are needed to differentiate between distinct species. The scientific community needs to be aware of this issue and needs to develop methods that better account for it when assigning taxonomic labels to metagenomic sequences from microbial communities.

## Main text

The sequencing of the 16S rRNA gene has commonly been employed to taxonomically characterize the prokaryotes found in microbial communities. Several 16S rRNA gene databases exist—RDP (1), Green Genes (2), SILVA (3)—comprising almost 4 million distinct gene variants. These databases are used as a reference when assigning taxonomic labels to newly sequenced versions of this gene. Although the 16S rRNA gene is universally present in the genomes of prokaryotes, and is phylogenetically-informative (i.e., useful for inferring their evolutionary history), its use to taxonomically-characterize microbial communities has been criticized due to the limited taxonomic resolution of 16S rRNA gene analyses (4, 5).

With the advent of metagenomics—the culture-independent sequencing and analysis of the total organismal DNA directly extracted from a sample—a broader range of methods have been developed to characterize the taxonomy of microbial communities. One commonality between metagenomic methods and those developed for the 16S rRNA is the use of reference databases to assign taxonomic labels to sequences. Instead of amplicon sequence variants (ASVs) or operational taxonomic units (OTUS), in a metagenomic context the database might consist of k-mers, genes, or genomes, and the sequences being taxonomically classified might be metagenomic reads, contigs, or metagenome-assembled-genomes (MAGs).

Here we demonstrate that the limited resolution of the 16S rRNA gene as a taxonomic marker reflects a more fundamental limitation of taxonomic classification itself, i.e., as sequence databases increase in size, the resolution of taxonomic classification made on the basis of these databases degrades due to the increased diversity of sequences with a specific taxonomic label. This effect has previously been demonstrated for whole-metagenome analyses using a k-mer classifier trained on various versions of the NCBI RefSeq database (6). Here, we demonstrate that the same pattern is observed when analyzing marker gene sequences, including the 16S rRNA gene and single copy marker genes that are phylogenetically informative and nearly universally present in prokaryotes (4, 7). The marker genes we analyzed included a set of 40 single copy marker genes (8), and the 120 genes used by the Genome Taxonomy Database (GTDB) (9) (these two sets overlap by 3 genes).

We began our analysis by comparing the effect of database size on the SILVA 16S rRNA gene database (394,617 sequences after removal of sequences with incomplete taxonomic labels as well as those from mitochondria and plastids) and the 120 marker genes used by the Genome Taxonomy Database (GTDB) project (35,171,383 total sequences). It should be noted that both databases contain full-length genes that have not been deduplicated, i.e., there can be identical sequences from the same species. To assess the level of ambiguity present in the database in a classifier-independent manner, we clustered the database sequences at different identity cut-offs and computed the number of clusters that contain sequences from more than one species (multi-species clusters). Multi-species clusters approximate the likelihood that a classifier would be unable to distinguish between the distinct species found in the cluster. It is also important to note that here we cluster the database sequences rather than the query sequences (as would normally be done if we constructed ASVs or OTUs). For each gene, we created a collection of random subsets varying in size from 10,000 to 220,000 sequences. Each subset was clustered with CD-Hit (10) at several sequence identity cut-offs (95%, 97%, 99%, 100%), requiring that shorter sequences fully align to longer ones. The clustering thresholds were chosen for several reasons. Firstly, genes sharing high sequence similarity are likely to contain regions where metagenomic reads cannot be uniquely aligned, thus, highlighting issues with read-based taxonomic classification. Further, genes from distinct species that are 100% identical indicate that species-level resolution is simply not possible for a specific gene. Secondly, these thresholds are often applied in various bioinformatic contexts. For example, microbial gene catalogs are often clustered at 95% identity to produce species-level gene clusters (11), while 97% and 99% identity have been used as proxy species-level thresholds in OTU clustering (12).

For all genes, the number of sequences in multi-species clusters increased at a super-linear rate as the database grew (Figures 1A and 1B), with higher rates of growth when clustered at higher similarity thresholds. The rate at which sequences are clustered with sequences from other species was estimated using the *Y = cX*^*m*^ linear regression model in log-log space, where *m* is the rate, *Y* is the number of sequences in multi-species clusters, and *X* is the number of genes in the simulated database. The fraction of species that had at least one sequence in a multi-species cluster increased with database size (Figure 1C); albeit this fraction was lower when clustering with higher similarity thresholds. The number of multi-species clusters depended on the density at which a particular taxonomic “neighborhood” was sampled, with a positive linear correlation between the number of distinct species within a genus and the number of multi-species clusters formed by sequences from that genus (Figure 1D). Importantly, amongst the species pairs that most frequently cooccurred in multispecies clusters when clustered at 100% identity were pathogens like *Bacillus anthracis* and *Vibrio cholerae*, which clustered with non-pathogenic species from their respective genera (Supplementary Table 1). This observation highlights the difficulty of distinguishing important human pathogens from their non-pathogenic neighbors as databases grow.

**Figure 1:**
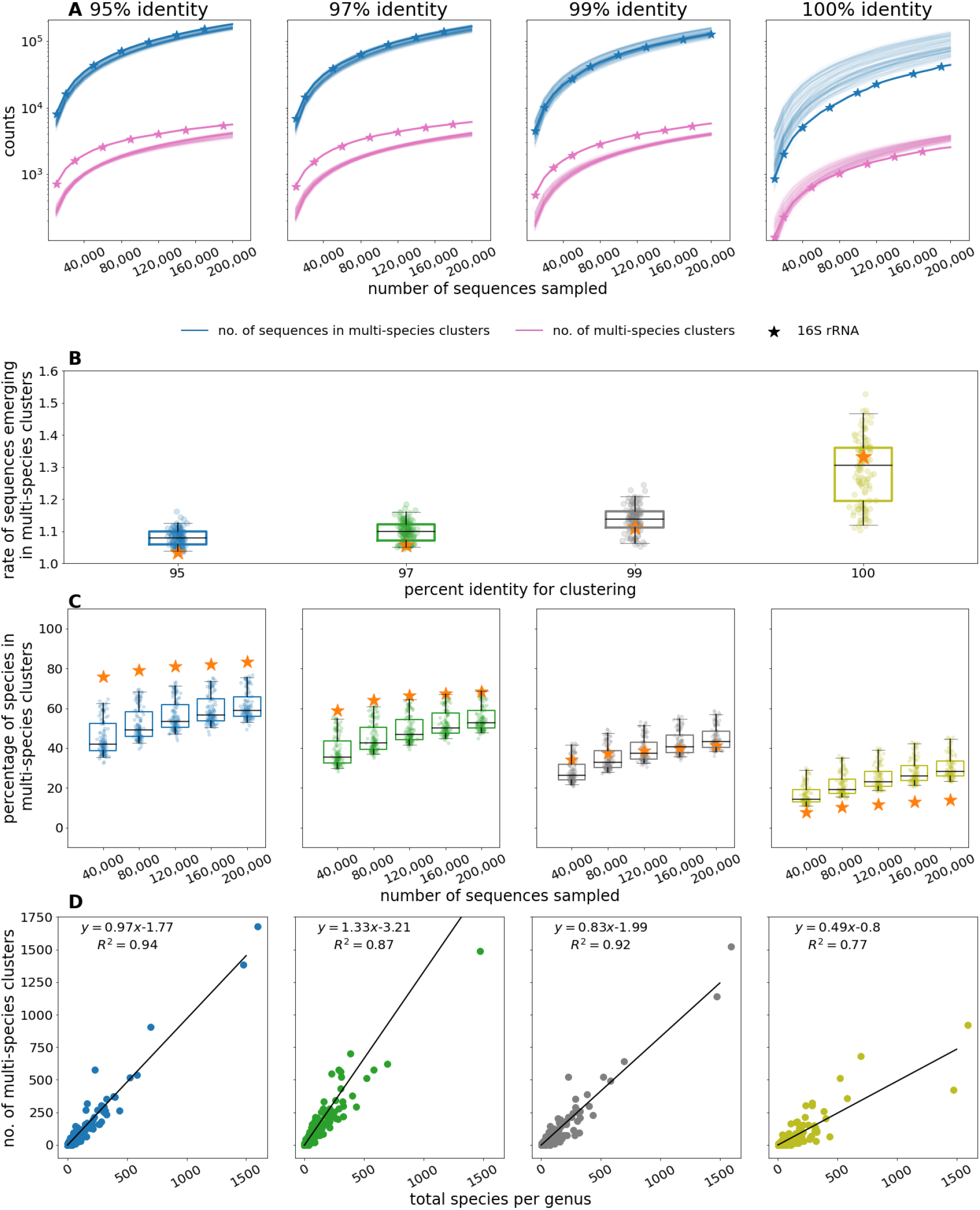
Clustering analysis for simulated databases created by randomly sampling sequences from the 16S rRNA SILVA database and the 120 marker gene Genome Taxonomy Database (GTDB). Each simulated database was clustered with CD-Hit at several sequence identity cut-offs (95%, 97%, 99%, 100%), requiring that shorter sequences fully align to longer ones. The 16S rRNA gene is denoted by a star in all subplots. A) The relationship between the number of genes in the simulated databases and the number of sequences in multi-species clusters and the total number of multi-species clusters. For the GTDB, each curve is for one of the 120 marker genes. B) The slopes of the regression models for the number of sequences in multi-species clusters as a function of the number of genes in the simulated databases. Each point represents one of the 120 marker genes in the GTDB. C) The proportion of species with sequences in multi-species clusters. D) The relationship between the number of multi-species clusters that a species belongs to and the species richness of its genus (i.e., the total number of species from that genus) in the simulated database. This data was only taken from the final iteration of the simulated databases. The results were aggregated across all 120 marker genes in the GTDB.

To further highlight the extent to which database growth impacts the specificity of taxonomic labels, we focused on a single genus, *Listeria*. This genus, that includes the important foodborne pathogen *Listeria monocytogenes*, comprised 5,014 RefSeq genome sequences, representing 34 distinct species. Most genome sequences from this genus (4,439) belonged to *Listeria monocytogenes*. From each *Listeria* genome sequence we extracted the 16S rRNA gene using Barrnap (13) (7,625 sequences) and 40 prokaryotic marker genes using fetchMG (14).

We identified 7,625 16S rRNA gene sequences (ranging from 1 to 9 copies per genome, consistent with the expectation that this gene is often multicopy in *Listeria*) and 200,359 marker gene sequences (~40 per genome, consistent with the expectation that these genes are single copy). The output of fetchMG was filtered, for each marker gene, by removing sequences that were of very different length (below half or above twice as long as the average sequence) and, thus, likely to be artifacts. For each gene, we randomly subsampled its sequences into sets of varying sizes, from 1,000 to 5,000 gene sequences in 1,000 gene increments. We repeated this process 100 times to estimate the variability of our results. As observed more broadly, even within this single genus, the number of sequences in multi-species clusters increased with the database size at all clustering thresholds and for all marker genes, including the 16S rRNA gene (Figure 2). Notably, at 95% identity each *Listeria* species had 16S rRNA gene sequences in multi-species clusters when analyzing the full data set. Further, 25% (100% identity) to 65% (95% identity) of all the *L. monocytogenes* sequences occurred in multi-species clusters, reiterating that database growth is affecting our ability to discriminate important pathogens from their near neighbors.

**Figure 2:**
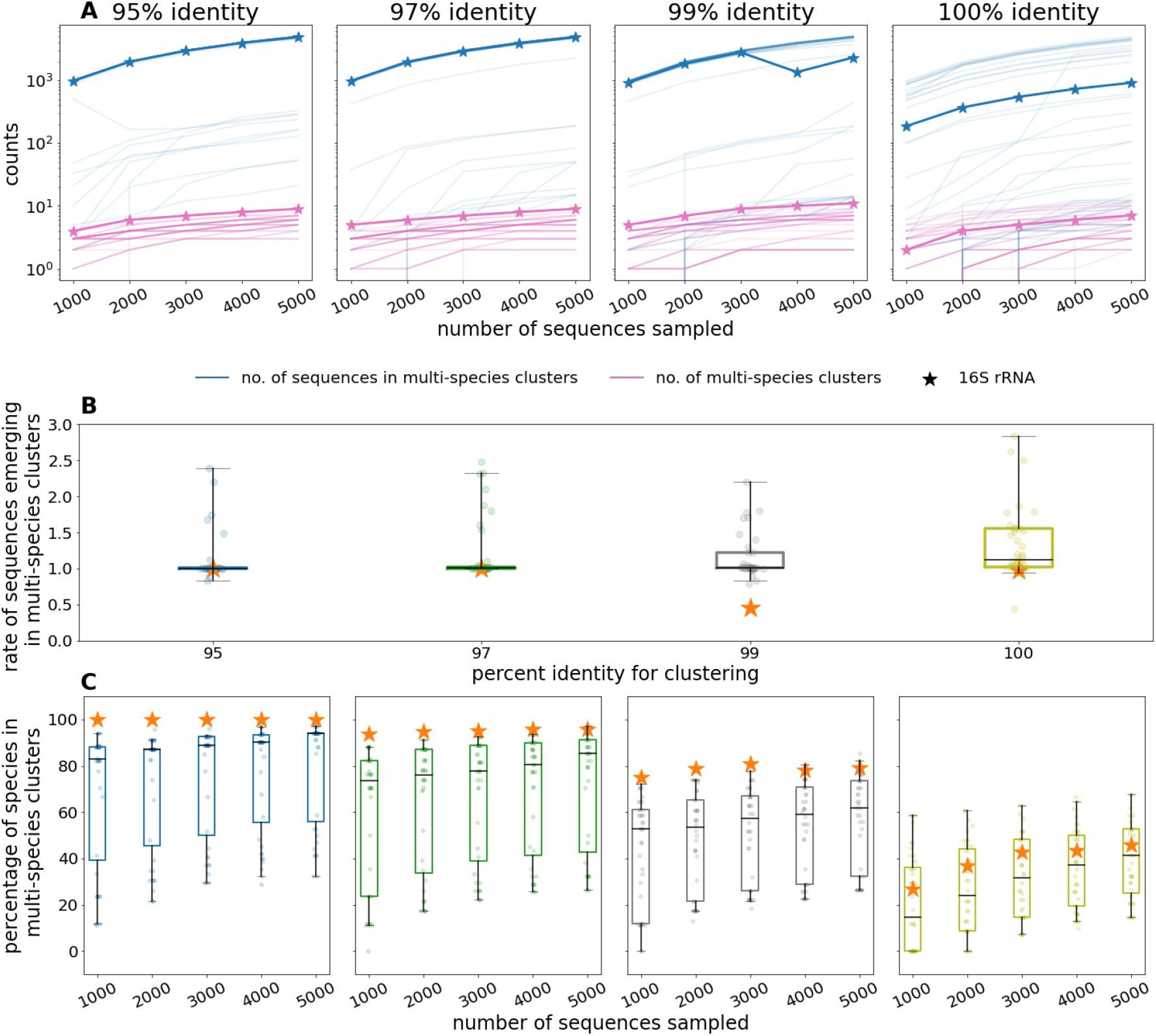
Clustering analysis for the simulated databases created by randomly sampling sequences from the 16S rRNA and the 40 marker genes extracted from 5,014 *Listeria* genomes. Each simulated database was clustered with CD-Hit at several sequence identity cut-offs (95%, 97%, 99%, 100%), requiring that shorter sequences fully align to longer ones. The results for each gene are reported by the median over 100 bootstrap experiments. The 16S rRNA gene is denoted by a star in all subplots. A) The relationship between the number of genes in the simulated databases and the number of sequences in multi-species clusters and the total number of multi-species clusters. Each curve is for one of the 40 marker genes B) The slopes of the regression models for the number of sequences in multi-species clusters as a function of the number of genes in the simulated databases Each point represents one of the 40 marker genes C) The proportion of species with sequences in multi-species clusters.

Our results support the observation that as sequence databases grow to contain more species and sequences, they are progressively losing species-level resolution for taxonomic classification (6). This observation has broad implications given that genes are routinely employed for various bioinformatic tasks, such as the taxonomic classification of genomic and metagenomic reads and assemblies, the curation of gene and metagenome-assembled-genome (MAG) catalogs, phylogeny estimation, and abundance profiling (3, 9, 11, 15, 16). The main difference between marker genes was the rate at which they lost species-level resolution, with the 16S rRNA gene sometimes being an outlier (a previously noted trend (4, 5)). Notably, at lower clustering thresholds (corresponding to longer evolutionary distances), the 16S rRNA gene is less informative than other marker genes, while at the highest threshold of 100% identity the 16S rRNA gene shows higher discrimination, likely due to the higher level of short-term sequence evolution within hypervariable regions of this gene.

Our results call into question highly specific taxonomic classifications (e.g., species, strain) for individual metagenomic reads and genes that are simplistically assigned based upon sequence similarity to database sequences. While, at some level, the community understands that accurate classification requires the use of whole-genome data (9, 17, 18), many computational tools for metagenomic analysis continue to classify individual k-mers, reads, and genes (1, 16, 19, 20). Our results indicate that such methods might provide overly specific taxonomic assignments for taxa that are underrepresented in the reference database used for analysis. Further, our results reemphasize the issues associated with the methods used to create microbial gene catalogs, which typically cluster taxonomically unlabeled genes into a set of representative sequences, which are then assigned taxonomic labels. Such workflows have been shown to create multispecies clusters, irrespective of the sequence identity threshold used for clustering, thereby effectively erasing entire species from the catalogs (11).

We suggest that taxonomic classifiers account for how densely a particular taxonomic group is represented in a database when assigning taxonomic labels. Further, future work should continue to explore what fraction of a genome is required to accurately identify a particular species within metagenomic data, and to quantify the taxonomic information content of different genomic regions. Instructively, even when comparing the average nucleotide identity (ANI) of the entire shared gene or genomic content of genomes, the commonly used threshold of 95% ANI is inconsistent as a proxy for species. For example, there are species within genera like *Brucella* or *Mycobacterium* that can only be differentiated above 99.5% ANI (18, 21). And highly sampled species (≥100 genome sequences) have been observed to comprise strains that are more divergent than the 95% cutoff (22).

At the broadest level, our work reiterates the long-noted issues with using discrete and definitive taxonomic labels for DNA sequences that are continually evolving (23). Recombination alone can render a genome a mosaic of lineages, with some horizontally transferred sequences potentially unrelated to any specific lineage. And the implications extend beyond taxonomic analysis to analyses that rely on taxonomy. For example, it is common to use taxonomic classifications based upon marker genes to infer the functional profile of microbial communities (24). A better approach would combine the information from taxonomic and functional markers in an application-specific manner. It is important that future work continue exploring the relationship between molecular evolution, taxonomy, function, the composition of sequence databases, and the fidelity of annotations when using reference databases as substrates for metagenomic analysis.

## Data availability

All the data used for this study is publicly available. The SILVA database used can be downloaded from https://www.arb-silva.de/fileadmin/silva_databases/current/Exports/SILVA_138.1_SSURef_tax_silva.fasta.gz.

The GTDB marker genes (release version 207) can be downloaded from https://data.gtdb.ecogenomic.org/releases/release207/207.0/genomic_files_all/bac120_marker_genes_all_r207.tar.gz. The NCBI Assembly database accessions for the *Listeria* assemblies can be found in Supplementary File 1.

## Acknowledgements

None.

## Contributions

All authors contributed equally to this study. MP conceived of the study. MP, TL, HSM, and SC designed the experiments. TL, HSM, and SC conducted the experiments and performed the data analysis. All authors contributed to writing the manuscript.

## Supplementary information

Supplementary File 1 is a list of the NCBI accessions for the RefSeq assemblies used for the *Listeria* analysis.

